# Oncogenic ETS factors expressed from fusion genes utilize EWSR1 to associate with the paraspeckle and regulate transcription

**DOI:** 10.64898/2026.10.12.758628

**Authors:** Renee A. Kinne, Peter C. Hollenhorst

## Abstract

ETS family transcription factors are aberrantly expressed in prostate cancer and Ewing sarcoma due to chromosomal rearrangements resulting in fusion genes, most commonly *TMPRSS2::ERG* and *EWSR1::FLI1*, respectively. The products of these gene fusions regulate transcription together with the intrinsically disordered protein EWSR1. However, the role of EWSR1 as a transcriptional co-regulator is not fully understood. EWSR1 can undergo liquid-liquid phase separation. We found that ERG could integrate into EWSR1 phase separated droplets. We then tested the role of the paraspeckle, a nuclear membraneless organelle known to regulate transcription and include EWSR1. Both ERG and EWSR1::FLI1 associated with the paraspeckle scaffold long non-coding RNA NEAT1_2. Disruption of paraspeckles in prostate cancer and Ewing sarcoma cell lines resulted in disruption of oncogenic phenotypes mimicking the loss of oncogenic ETS factors. In Ewing sarcoma cells paraspeckle disruption promoted similar transcriptional profiles as loss of EWSR1::FLI1. Further, loss of EWSR1::FLI1 reduced the association of a paraspeckle component with EWSR1::FLI1 target genomic loci. In *TMPRSS2::ERG* positive prostate cancer cells paraspeckle disruption correlated with ERG-mediated transcriptional repression. These findings suggest that the paraspeckle is an important transcriptional regulator in cancers with ETS gene fusions and provides a potential target for further mechanistic understanding and drug development.

## Introduction

The ETS family of transcription factors is comprised of 28 members in humans. Genes encoding six ETS factors comprising the ERG (ERG, FLI1, FEV) and PEA3 (ETV1, ETV4, ETV5) subfamilies are altered by chromosomal rearrangement in prostate cancer and/or Ewing sarcoma. In prostate cancer, chromosomal rearrangements fuse the promoter and 5’ UTR of genes expressed in prostate epithelial cells to the open reading frame of *ERG, ETV1, ETV4,* or *ETV5* resulting in expression of either a full-length, or N-terminally truncated, ETS factor in a cell type where it is normally silent (1, 2). In Ewing sarcoma, rearrangements create a gene encoding a fusion protein with the N-terminus of EWSR1 (or EWSR1 homolog FUS) and the C-terminus (including DNA binding domain) of an ETS factor (FLI1, ERG, FEV, ETV1, or ETV4) (3). Although prostate cancer rearrangements usually do not lead to a fusion protein, we have previously found that the ETS factors rearranged in prostate cancer (ERG, ETV1, ETV4, and ETV5) bind to the EWSR1 protein (4), linking the function of EWSR1 to the gene fusion product transcription factor in both diseases. However, the mechanism through which EWSR1 affects ETS transcription factor function in these cancers has yet to be elucidated.

EWSR1 is an intrinsically disordered RNA binding protein (ID-RBP). The N-terminal ID region of EWSR1 promotes self-binding, forming phase separated granules, and can act as a transcriptional activation domain (5). Liquid-liquid phase-separation (LLPS) is a biochemical process that allows proteins to create transient and dynamic interactions. The process of phase separation has been linked with both tumorigenesis and general transcriptional regulation (6–8). The process of phase separation has been shown to be critical to the function of several different oncogenic transcription factor fusions (6). Therefore, the ability for EWSR1 to phase separate in the context of both prostate and Ewing sarcoma fusion transcription factors presents an interesting mechanism for exploration.

One potential mechanism of EWSR1 transcriptional co-regulation may be through the involvement of the paraspeckle, a phase separated membraneless organelle. The paraspeckle is a nuclear body with an organized ribonucleoprotein structure centered around the long non-coding RNA (lncRNA) NEAT1_2 (9). The NEAT1 gene encodes both long and short isoforms. The long isoform NEAT1_2 is required for paraspeckle formation and includes hairpin-like structures that bind critical paraspeckle proteins such as FUS and SFPQ/NONO heterodimers (10, 11). The short NEAT1 isoform, NEAT1_1 may localize to paraspeckles, but is not an essential component (12, 13). Paraspeckles have known roles in transcriptional regulation, primarily acting as a sequestering agent, and interact with open and active chromatin (14–16). In cancer, the paraspeckle has been shown to have both tumor suppressive and oncogenic roles, depending on the cancer background (14, 17–19).

There are several connections between transcription factors expressed due to gene fusions and paraspeckles. Ewing sarcoma fusion partner EWSR1 is found in paraspeckles, and FUS, a close homolog of EWSR1 that is sometimes fused to ETS in Ewing sarcoma, is a necessary paraspeckle component (20). NEAT1, the lncRNA that acts as the paraspeckle scaffold, has increased expression in prostate cancer patients (21). A study has also shown that NEAT1 can promote oncogenic phenotypes in VCaP cells, a prostate cancer cell line with a *TMPRSS2::ERG* fusion (14). Finally, SFPQ and NONO, two necessary paraspeckle components, are found fused to transcription factors such as TFE3 in translocation renal cell carcinoma, and SFPQ::TFE3 displays punctate distribution in cell nuclei (22). Identifying the paraspeckle as a mechanism of oncogenic EWSR1 co-activation in Ewing sarcoma and prostate cancer would further our understanding of the role of nuclear bodies in cancer and open new targets for drug development.

This study elucidated the role of paraspeckles in cancers dependent on ETS fusion genes. We found that ERG can integrate into EWSR1 phase separated droplets *in vitro*. We identified interactions between both ERG and EWSR1::FLI1 with the paraspeckle scaffolding lncRNA NEAT1_2 that were dependent on EWSR1. Disruption of paraspeckles in Ewing sarcoma cells inhibited colony formation and promoted migration, consistent with EWSR1::FLI1 loss. Loss of paraspeckles similarly inhibited colony formation in VCaP cells, a *TMPRSS2::ERG* positive prostate cancer line, showing that paraspeckle function is important for multiple ETS-positive cancer types. Transcriptome analysis showed that paraspeckle loss has a similar effect on gene expression as EWSR1::FLI1 loss in Ewing sarcoma and loss of ERG repression in prostate cancer. Finally, we identified paraspeckles at the chromatin of EWSR1::FLI1 direct targets, with this presence decreased with EWSR1::FLI1 knockdown. Together, these data support a model of oncogenic ETS modulating paraspeckle genomic loci to promote an oncogenic transcriptome.

## Results

### ERG participates in EWSR1 phase separation

The N-terminus of EWSR1 can promote phase separation (8, 23) and is the region of EWSR1 that directly binds ERG (24). Substitution of 37 tyrosine residues in the EWSR1 N-terminus with serine (YS37) disrupts phase separation (8) and also disrupts the interaction of EWSR1 with ERG (24). Therefore, we questioned whether these dual functions of the EWSR1 N-terminus; phase separation and ERG binding, can occur simultaneously or are mutually exclusive. To test this, droplet assays were performed using fluorescently tagged, purified, EWSR1 and ERG. LLPS was induced through the addition of 10% polyethylene glycol (PEG). EWSR1-mCherry formed droplets upon the addition of 10% PEG, but ERG-GFP did not (Figure 1A). To ensure this was LLPS and not protein aggregation, droplets were treated with either 1,6 hexanediol (1,6 HD), which is a disruptor of LLPS, or 2,5 hexanediol (2,5 HD), which is not a disruptor of LLPS (25, 26). EWSR1-mCherry droplet formation was disrupted by 1,6 HD but not 2,5 HD (Figure S1). While ERG was unable to undergo LLPS alone, when EWSR1 and ERG were combined before phase separation was induced, ERG formed droplets that co-localized with EWSR1 droplets (Figure 1B), indicating that ERG can participate in EWSR1 LLPS.

**Figure 1.**
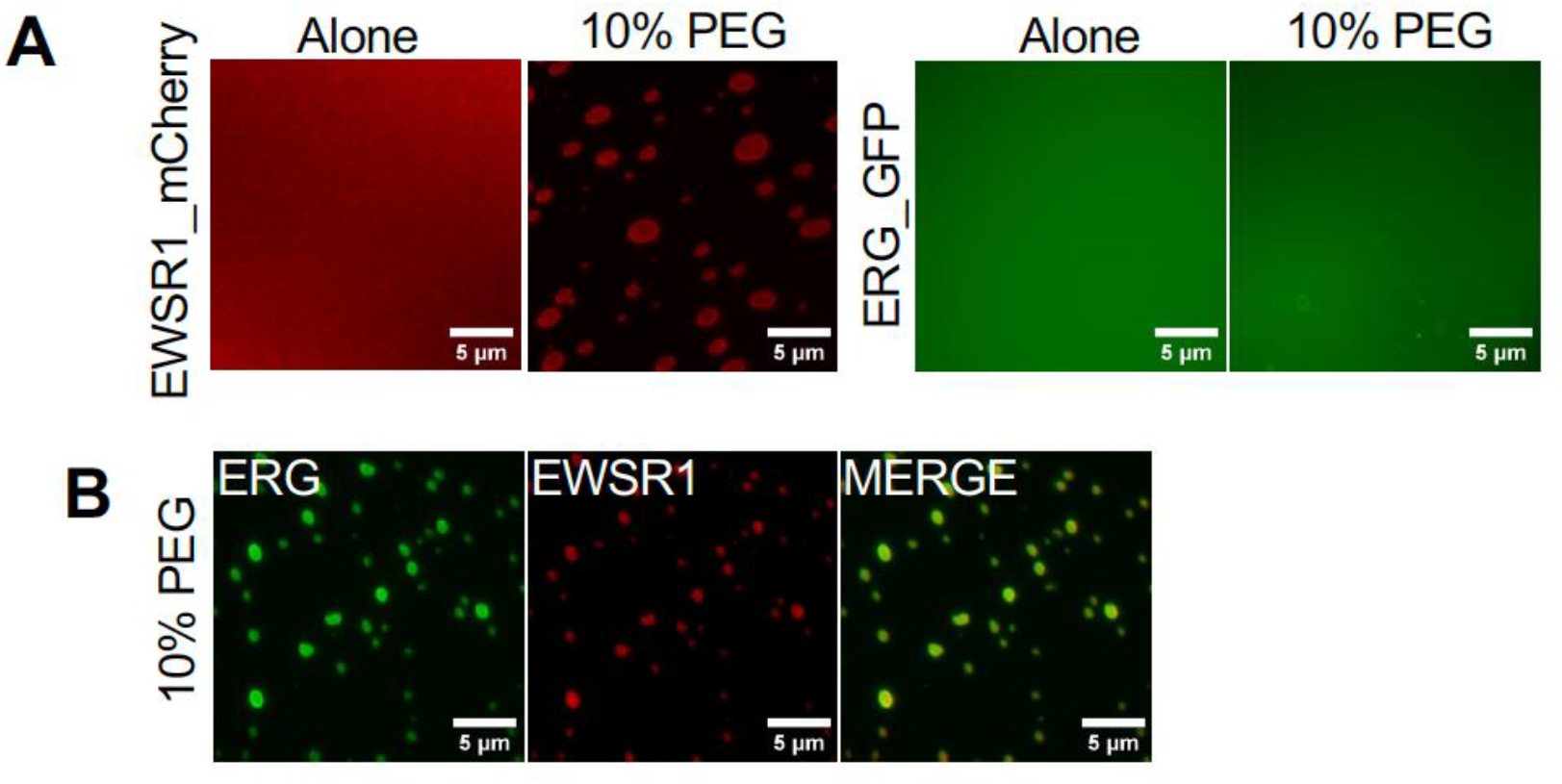
ERG participates in EWS phase separation. (A) Droplet formation assay using purified EWSR1_mCherry or ERG_eGFP with 10% PEG as a crowding agent to induce droplet formation. (B) EWSR1_mCherry and ERG_eGFP combined before 10% PEG addition. All proteins are at 3 µM starting concentration.

### NEAT1_2 interacts with oncogenic ETS through EWSR1 and forms nuclear puncti in VCaP cells

A potential mechanism of LLPS based regulation in ETS positive cancers is the involvement of the paraspeckle, a phase separated nuclear body. Paraspeckles are organized by the lncRNA NEAT1_2. RNA immunoprecipitation (RIP) of 3xFlag-EWSR1::FLI1 or 3xFlag-ERG exogenously expressed in HEK293t cells both showed enrichment of NEAT1_2 compared to empty vector pull down (Figure 2A). Because EWSR1 is found in paraspeckles, we predicted that the oncogenic ETS interaction with the paraspeckle would be dependent on EWSR1. ERG P436A, a mutant that disrupts ERG-EWSR1interaction (27), expressed at a similar level to ERG, did not show enrichment of NEAT1_2 compared to empty vector (Figure 1A). Further, expression of just the FLI1 portion of the EWSR1::FLI1 fusion also failed to enrich NEAT1_2. These findings support EWSR1 as a necessary connection between oncogenic ETS and NEAT1_2. To confirm that paraspeckles were present in cell lines expressing oncogenic ETS, we performed RNA-FISH for NEAT1 and immunofluorescence for SFPQ in VCaP cells, a *TMPRSS2::ERG* positive prostate cancer line. In VCaP cells NEAT1 and SFPQ, a necessary paraspeckle protein component, formed clear co-localized puncti which supports paraspeckle presence (Figure 2B). Co-staining for NEAT1 and ERG in VCaP cells showed that NEAT1 and ERG were found in overlapping nuclear regions, but ERG was not uniquely sequestered in paraspeckles (Figure 2C).

**Figure 2.**
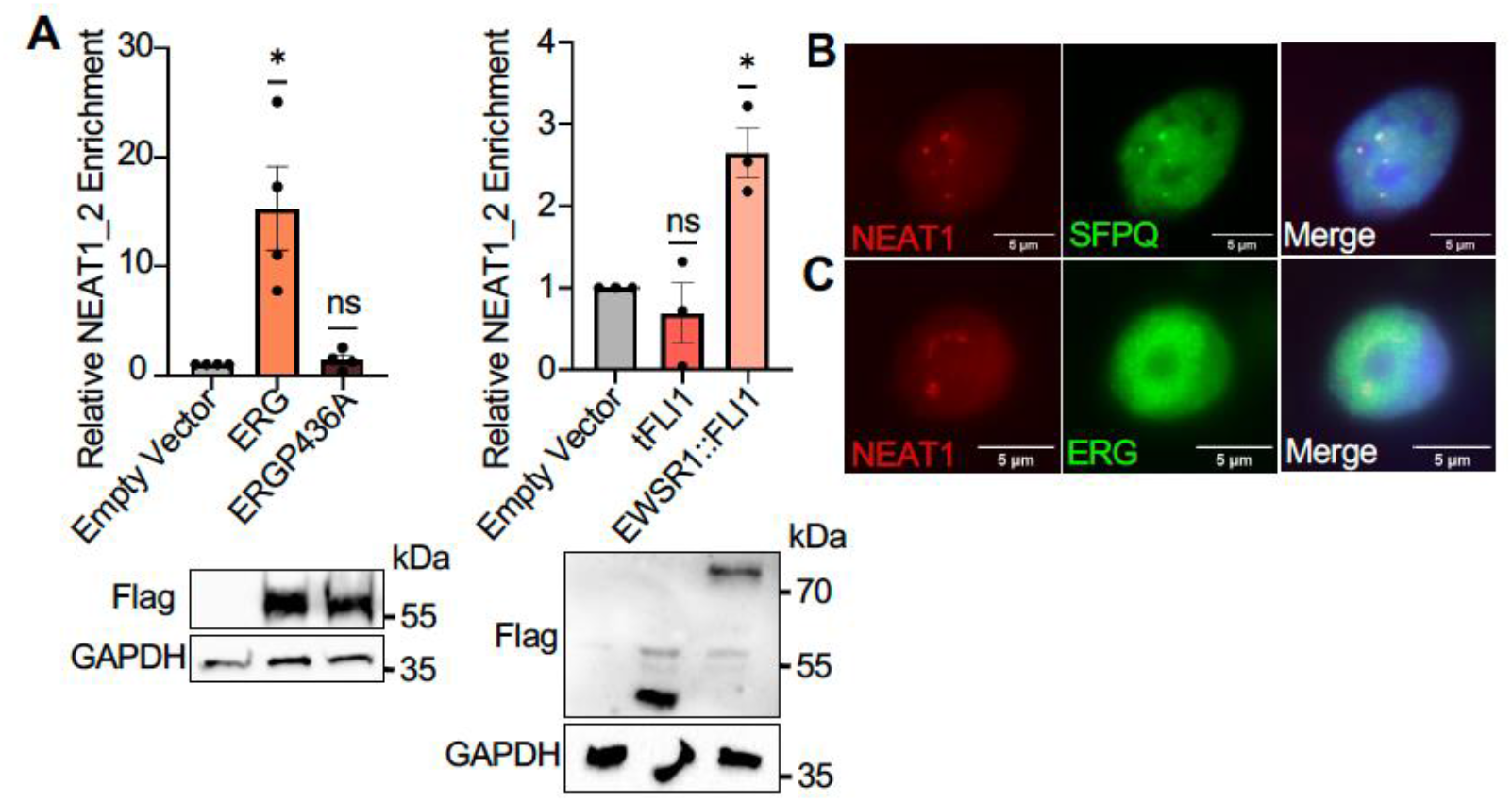
NEAT1_2 interacts with oncogenic ETS through EWSR1 and forms nuclear puncti in VCaP cells. (A) RNA immunoprecipitation (RIP) with Flag antibody of 3xFlag tagged ETS or empty vector. Reverse transcription quantitative PCR (RT-qPCR) measured NEAT1_2 and normalized to empty vector. ERG RIPs n=4; EWSR1::FLI1 RIPs n=3. Mean and SEM shown. Statistical significance was calculated by one sample t-test *p < 0.05. Immunoblot showing relative expression of Flag-tagged proteins (lower). (B) Immunofluorescence and in situ hybridization (IF-FISH) in VCaP cells. NEAT1: red SFPQ: green DAPI: blue. C. IF-FISH in VCaP cells. NEAT1: red; ERG: green; DAPI: blue.

### Paraspeckle disruption inhibits EWSR1::FLI1 dependent phenotypes in Ewing sarcoma cells

NEAT1_2 and FUS are both necessary paraspeckle components, with knockdown of these components disrupting paraspeckle formation (9, 10, 28). To determine if the paraspeckle has a role in the oncogenic function of EWSR1::FLI1, NEAT1 and FUS were targeted for knockdown in A673 Ewing sarcoma cells. Two independent short hairpin RNA (shRNA) were developed for each target (Figure 3A and B). For NEAT1 knockdown one shRNA targeted the long isoform that acts as the paraspeckle scaffold, denoted as NEAT1_2, and one targeted both the long and short isoforms denoted as NEAT1. Knockdown of NEAT1 reduced NEAT1 FISH signal and disrupted NEAT1 nuclear puncti formation (Figure 3C). Knockdown of FUS disrupted NEAT1 nuclear puncti, but also redistributed NEAT1 to the cytoplasm, where it formed new cytoplasmic puncti (Figure 3C). Therefore, both NEAT1 and FUS knockdown disrupted nuclear paraspeckles. A673 cells require EWSR1::FLI1 for anchorage independent growth, but not normal proliferation on plastic (29). Knockdown of both FUS and NEAT1 reduced anchorage independent growth in a soft agar assay (Figure 3D) but did not reduce proliferation on plastic (Figure 3F), consistent with loss of EWSR1::FLI1 function (Figure 3E and F). EWSR1::FLI1 is known to reduce mesenchymal cell differentiation in Ewing sarcoma, with knockdown of EWSR1::FLI1 leading to a more differentiated mesenchymal state and an increase in cell migration in A673 (29, 30). We found that EWSR1::FLI1 knockdown increased A673 migration more than 10-fold (Figure 3G). Similarly, knockdown of both NEAT1 and FUS increased migration of A673 cells (Figure 3G). Unlike A673 cells, TC71 Ewing sarcoma cells require EWSR1::FLI1 for normal proliferation on plastic. Consistent with this, introduction of an shRNA targeting EWSR1::FLI1 to TC71 cells reduced proliferation as compared to a negative control shRNA targeting luciferase (Figure 3H). Similarly, introduction of shRNAs targeting NEAT1 and FUS reduced TC71 proliferation to the same extent as targeting EWSR1::FLI1 (Figure 3H). This is in contrast to A673 cells where these same shRNA constructs have no effect on proliferation (Figure 3F). Together, these data support the hypothesis that paraspeckles are necessary for EWSR1::FLI1 function in Ewing sarcoma cells.

**Figure 3.**
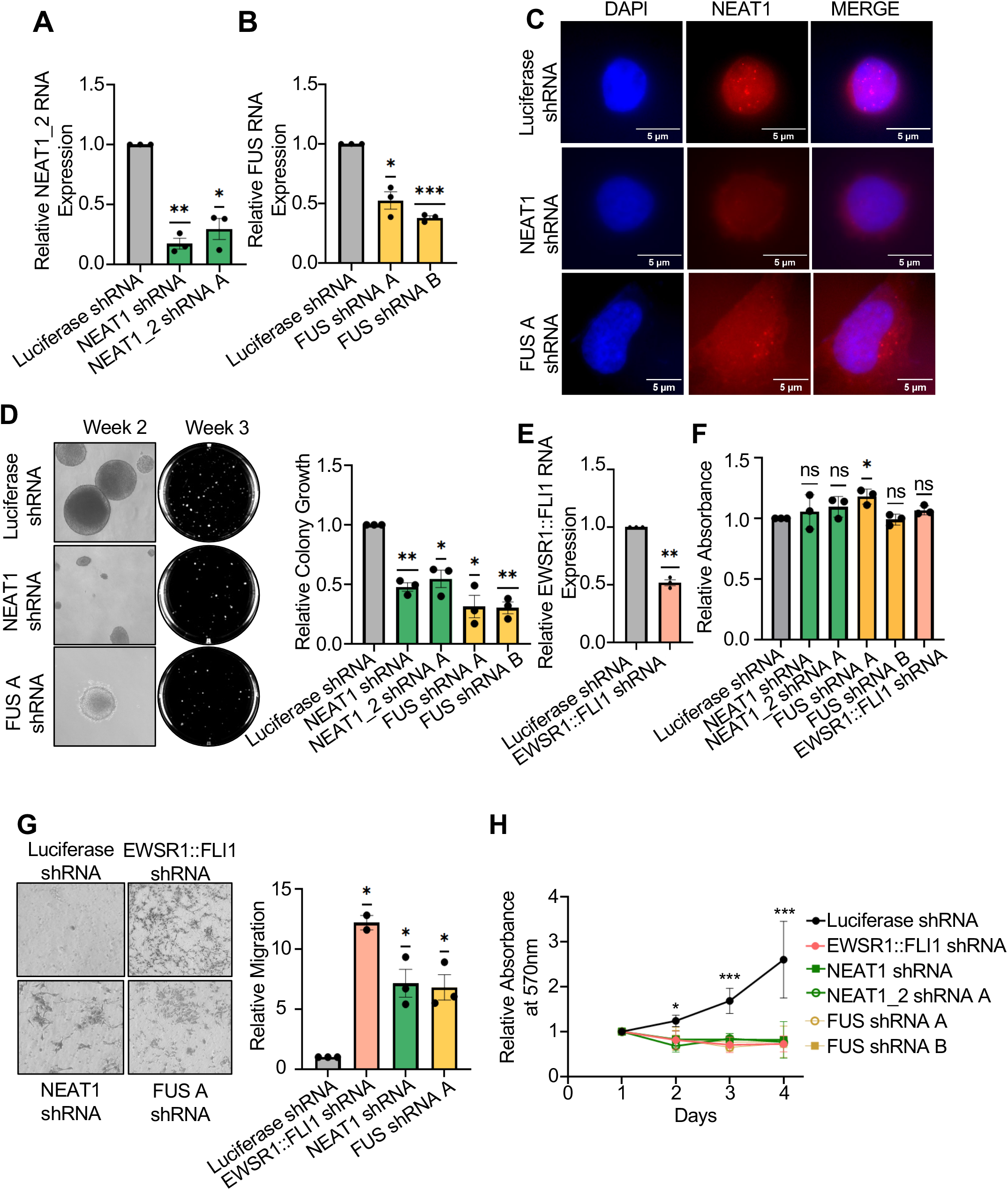
Disruption of paraspeckles in A673 cells disrupts oncogenic phenotypes. (A) RT-qPCR quantification of NEAT1_2 RNA in A673 cells with shRNA targeting luciferase, NEAT1, or NEAT1_2 normalized first to housekeeping control EF1α and then to negative control luciferase shRNA (B) RT-qPCR quantification of FUS RNA in A673 with shRNA targeting luciferase or FUS normalized to EF1α then to luciferase shRNA. (C) NEAT1 RNA-FISH in A673 cells with luciferase shRNA, NEAT1 shRNA, or FUS shRNA. NEAT1: Red; DAPI: blue. (D) Soft agar assay of A673 cells with luciferase shRNA, NEAT1 shRNA, NEAT1_2 shRNA or FUS shRNA. Representative microscopic images of colonies were taken at 2 weeks. Images of entire wells were taken at 3 weeks. Area of colony growth was quantified at 3 weeks and normalized to luciferase shRNA. (E) RT-qPCR quantification of EWSR1::FLI1 RNA in A673 cells with shRNA targeting luciferase or EWSR1::FLI1 normalized to EF1α then to luciferase shRNA (F) MTT assay for proliferation in A673 cells with shRNA targeting luciferase, NEAT1, NEAT1_2, or FUS. Day 3 absorbance was normalized to day 1 of same shRNA and then to luciferase shRNA. (G) Trans well migration assay of A673 cells with shRNA targeting luciferase, NEAT1, FUS, or EWSR1::FLI1. Representative images shown. Area of cell growth on membrane was normalized to luciferase shRNA. In EWSR1::FLI1 shRNA error bars are SEM of 2 biological replicates. (G) MTT assay for proliferation of TC71 cells with shRNA targeting luciferase, NEAT1, NEAT1_2, or FUS. Absorbance for each day was normalized to day 1 of same shRNA. *p*-values were calculated using one way ANOVA, where ∗ indicates a *p*-value < 0.05 and ∗∗∗ indicates a *p*-value < 0.001 Unless stated otherwise, error bars are SEM of three biological replicates. Unless stated otherwise *p*-values were calculated using one sample *t* test, where ∗ indicates a *p*-value < 0.05, ∗∗ indicates a *p*-value < 0.01, and ∗∗∗ indicates a *p*-value < 0.001

### Paraspeckle oncogenic phenotypes in VCaP cells

To determine if paraspeckles have a role in ERG function in prostate cancer cells, NEAT1 and FUS were knocked down in VCaP cells, a *TMPRSS2::ERG* positive prostate cancer cell line (Figure 4A and B). Notably, NEAT1 shRNA knockdown in VCaP cells was not as robust as in A673 cells. Consistent with this, knockdown of NEAT1 did not reduce FISH signal to the same degree as in A673 cells, however puncti observed in the scramble shRNA control were lost (Figure 4C). Similar to the A673 cells, FUS shRNA in VCaP resulted in loss of nuclear NEAT1 puncti but gain of cytoplasmic puncti (Figure 4C). ShRNA knockdown of ERG in VCaP cells led to a reduction in colony formation (Figure 4D-E, and S2). Knockdown of both NEAT1 and FUS in VCaP cells also significantly decreased colony formation (Figure 4E). Proliferation was not significantly reduced with paraspeckle disruption (Figure 4F). These results are consistent with our A673 model and similar to the effects of EWSR1 knockdown in VCaP cells (4, 24). Together, these data support a hypothesis that the paraspeckle plays a role in ERG function in VCaP cells.

**Figure 4.**
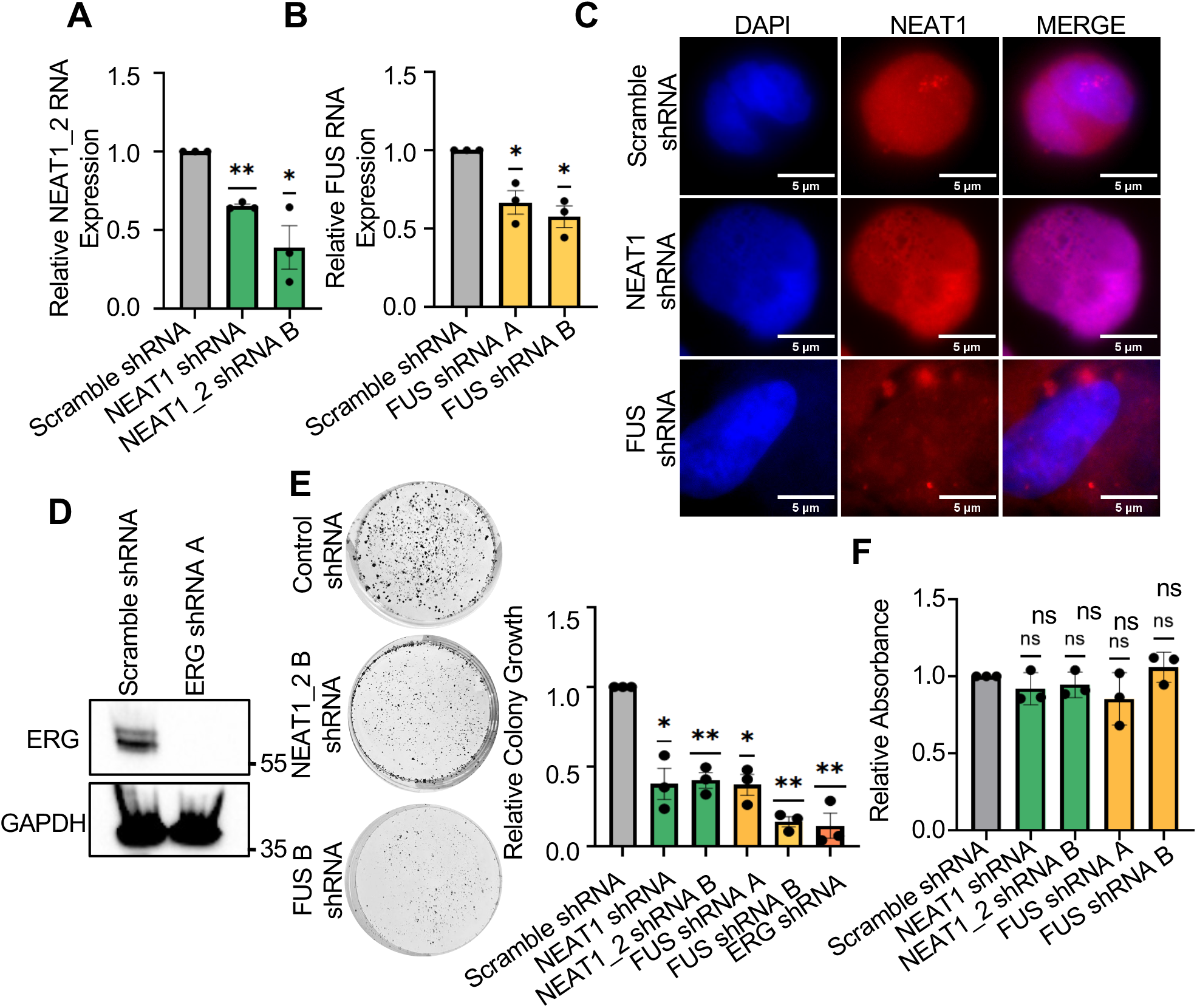
Paraspeckle disruption in VCaP cells inhibits colony formation. (A) RT-qPCR quantification of NEAT1_2 RNA in VCaP with scramble shRNA, or shRNA targeting NEAT1 or NEAT1_2 normalized first to EF1α and then to scramble shRNA. (B) RT-qPCR quantification of FUS RNA in VCaP scramble shRNA or FUS shRNA cells normalized as in (A). (C) NEAT1 RNA-FISH in VCaP cells with scramble shRNA, or shRNA targeting NEAT1, or FUS. NEAT1: Red; DAPI: blue. (D) Immunoblot with ERG or GAPDH antibody of VCaP cell lysate with either scramble shRNA or ERG shRNA. (E) Colony formation assay of VCaP cells with scramble shRNA or shRNA targeting NEAT1, NEAT1_2, FUS, or ERG. Area of colony growth was quantified at 3 weeks and normalized to scramble shRNA. Representative images on left. ERG shRNA colony formation is represented by the average of single biological trials using 3 different shRNAs targeting ERG as shown in (D) and Fig S2. (F) MTT assay for proliferation in VCaP cells with scramble shRNA, or shRNA targeting NEAT1, NEAT1_2, or FUS. MTT absorbance on day 3 was normalized to day 1 and then normalized to scramble shRNA. Error bars are SEM of three biological replicates. *p*-values were calculated using one sample *t* test, where ∗ indicates a *p*-value < 0.05, ∗∗ indicates a *p*-value < 0.01

### EWSR1::FLI1 associates paraspeckles with target genes to regulate transcription

The paraspeckle has been reported to have roles in regulating transcription (14, 15). To test if paraspeckle disruption can alter the transcriptional regulatory role of EWSR1::FLI1, RNA-seq was performed in A673 cells that expressed shRNAs targeting luciferase (control), EWSR1::FLI1, NEAT1, or FUS. The differentially expressed genes (DEGs) in both NEAT1 and FUS knockdown significantly overlapped, with about 30% of the DEGs up or downregulated in FUS knockdown also showing that same change in NEAT1 knockdown (Figure 5A). When all EWSR1::FLI1 DEGs were aligned with NEAT1 and FUS knockdown, there were strikingly similar trends between all three groups (Figure 5B). Gene set enrichment analysis (GSEA) of EWSR1::FLI1, NEAT1, and FUS knockdown showed similar decreases in pathways associated with pro-growth transcription factors E2F and MYC (Figure S2). There was also correlation for all 3 data sets with upregulation of epithelial to mesenchymal transition (Figure 5C) that is consistent with gain of cellular migration (Figure 3G). Together, these data suggest that the paraspeckle and EWSR1::FLI1 together promote loss of mesenchymal differentiation within Ewing sarcoma cells.

**Figure 5.**
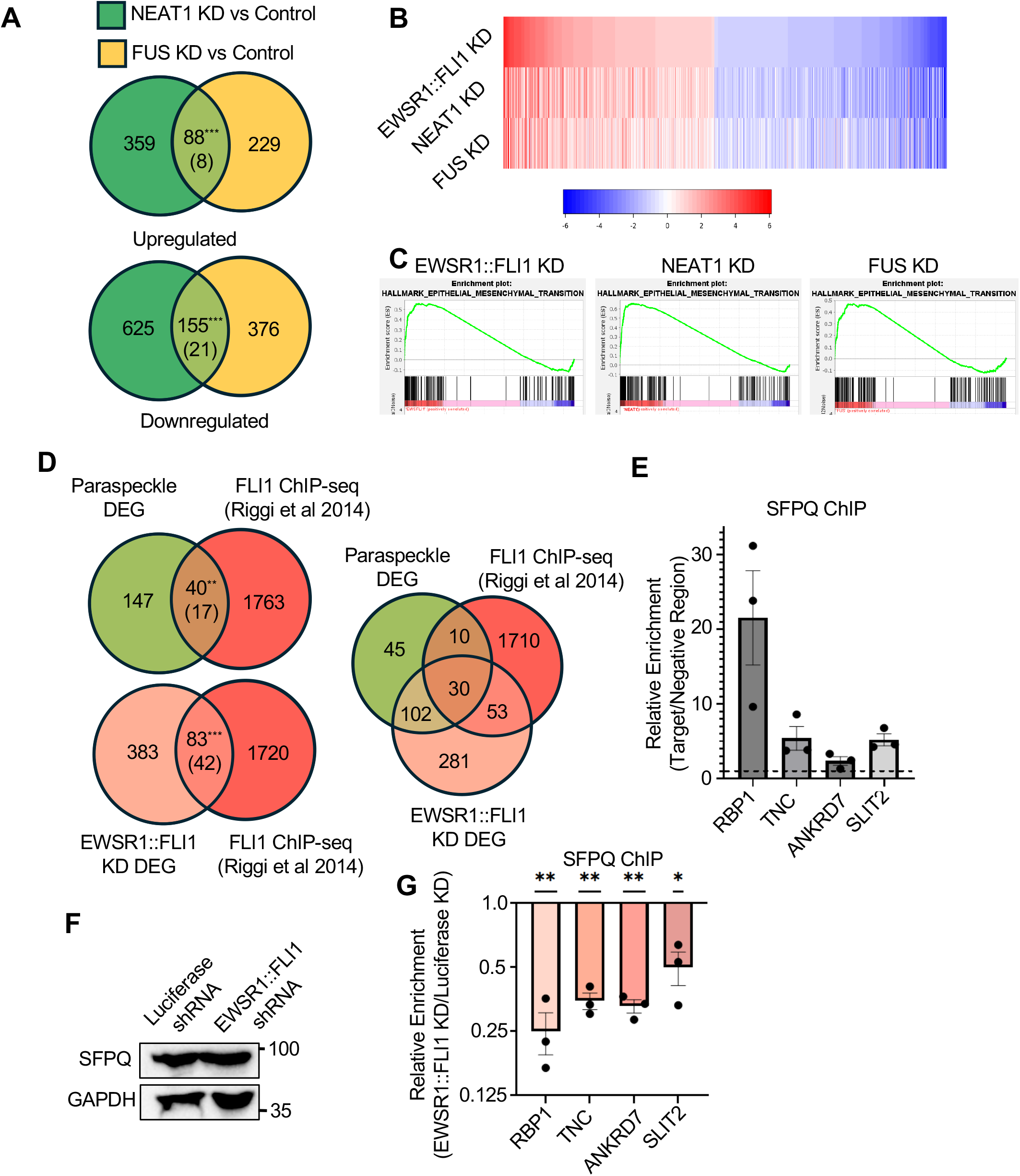
Paraspeckles regulate EWSR1::FLI1 transcription. (A) Overlap of differentially upregulated or downregulated genes in NEAT1 or FUS shRNA compared to luciferase shRNA (control) in A673 cells. Number in parenthesis represents random expected overlap. *p*-values were calculated using fishers exact test, where ∗∗∗ indicates a *p*-value < 0.001 (B) Heat map comparison of EWSR1::FLI1 shRNA, NEAT1 shRNA, or FUS shRNA each compared to luciferase shRNA in A673 cells. Genes shown were identified and ordered based on EWSR1::FLI1 differentially expressed genes (DEG) with padj<0.05 and |log(FC)|=> 1 cut off. (C) Gene set enrichment analysis compared to hallmark epithelial to mesenchymal transition genes for EWSR1::FLI1 shRNA, NEAT1 shRNA, and FUS shRNA each in comparison to luciferase shRNA. (D) Overlap of DEGs in both FUS shRNA and NEAT1 shRNA (paraspeckle) compared to nearest gene in FLI1 ChIP-seq from A673 cells (upper left) or overlap of EWSR1::FLI1 DEGs compared to FLI1 ChIP-seq (lower left). Right side shows overlap of all 3 groups. Number in parenthesis represents random expected overlap. *p*-values were calculated using fishers exact test, where ∗∗ indicates a *p*-value < 0.01 and ∗∗∗ indicates a *p*-value < 0.001 (E) ChIP-qPCR of SFPQ at previously established EWSR1::FLI1 bound regions with in A673 cells expressing luciferase shRNA. Enrichment was determined by normalization to a genomic region not known to be bound by EWSR1::FLI1. Dashed line at 1-fold. (F) Immunoblot for SFPQ or GAPDH in A673 cells with luciferase shRNA or EWSR1::FLI1 shRNA. (G) Ratio of SFPQ ChIP-qPCR enrichment at the same targets as in (E) comparing A673 cells EWSR1::FLI1 shRNA to A673 cells expressing luciferase shRNA. Error bars are SEM of three biological replicates. *p*-values were calculated using one sample *t* test, where ∗ indicates a *p*-value < 0.05, ∗∗ indicates a *p*-value < 0.01.

To test if genes regulated by the paraspeckle correlated with direct EWSR1::FLI1 target genes, RNA-seq was compared to published ChIP-seq mapping data. Genes differentially expressed in EWSR1::FLI1 knockdown were enriched when compared to a list of nearest genes to FLI1 ChIP-seq peaks in A673 cells (Figure 5D, lower). All DEGs that matched in NEAT1 and FUS knockdown (Figure 5A) were then compiled into a paraspeckle signature of 243 genes. Paraspeckle signature genes were also significantly enriched when compared to the nearest gene to FLI1 ChIP-seq peaks in A673 cells (Figure 5C, upper) (31). Of the 40 genes that were found in both the paraspeckle signature and FLI1 ChIP-seq, 30 were also differentially expressed with EWSR1::FLI1 knockdown (Figure 5C, right). This indicates that the paraspeckle can participate in the transcriptional regulation of direct EWSR1::FLI1 transcriptional targets. Interestingly, we did not observe any difference in overlap between upregulated and downregulated genes, suggesting a role in both processes. It has been previously published that paraspeckles can dock directly at transcriptional start sites to regulate gene expression (32). We hypothesized that EWSR1::FLI1 can direct paraspeckles to specific genomic loci to promote regulation of target genes. To test this, candidate regulatory regions from the overlap of the FLI1 ChIP-seq and the paraspeckle signature were tested for enrichment in ChIP-qPCR for paraspeckle component SFPQ in A673 cells. These candidates included genes that were both activated and repressed during paraspeckle disruption. SFPQ enriched all of the tested genomic loci, suggesting proximal location of paraspeckles (Figure 5E). SFPQ protein levels were not affected by EWSR1::FLI1 knockdown (Figure 5F). SFPQ ChIP-qPCR was repeated in EWSR1::FLI1 knockdown A673 cells. ChIP enrichment was significantly reduced with EWSR1::FLI1 knockdown in every target tested (Figure 5G) indicating that SFPQ enrichment at these loci is dependent on EWSR1::FLI1. These data support a model that EWSR1::FLI1 directs the paraspeckle to genomic targets to regulate transcription.

### Paraspeckles mediate ERG repression in prostate cancer cells

Considering the similarity in function of ERG and EWSR1::FLI1 and the consistency of phenotypic responses to paraspeckle disruption in VCaP and A673 cells, we hypothesized that paraspeckles regulate transcription through a similar mechanism in both Ewing sarcoma and ERG positive prostate cancer. To test this, RNA-seq measured gene expression in VCaP prostate cancer cells with either control, NEAT1, or FUS knockdown. There was a significant overlap in both upregulated and downregulated genes between NEAT1 and FUS knockdown (Figure 6A). There was very little overlap between the paraspeckle signature in A673 cells and the 749 DEGs in common between NEAT1 and FUS knockdown in VCaP cells with only 10 genes on both lists (Figure S3A). Interestingly, when gene ontology enrichment analysis of the upregulated paraspeckle signature in VCaP cells identified multiple ERG-related functions (Figure 6B). ERG is canonically expressed in hematopoietic stem cells with roles in stem cell maintenance (33, 34). The genes upregulated when paraspeckles were knocked down were enriched for functions involving a loss of stemness, such as activation of epithelial differentiation and cell fate commitment (Figure 6B). This connection was not observed in the downregulated paraspeckle signature (Figure S3B). When compared to publicly available ERG knockdown RNA-seq in VCaP cells (35), genes in the upregulated paraspeckle signature were significantly correlated with increased expression during ERG knockdown (Figure 6C). However, genes in the downregulated paraspeckle signature were not correlated with increased or decreased expression with ERG knockdown (Figure 6C). Together, these data are indicative of paraspeckles playing a role in ERG mediated transcriptional repression.

**Figure 6.**
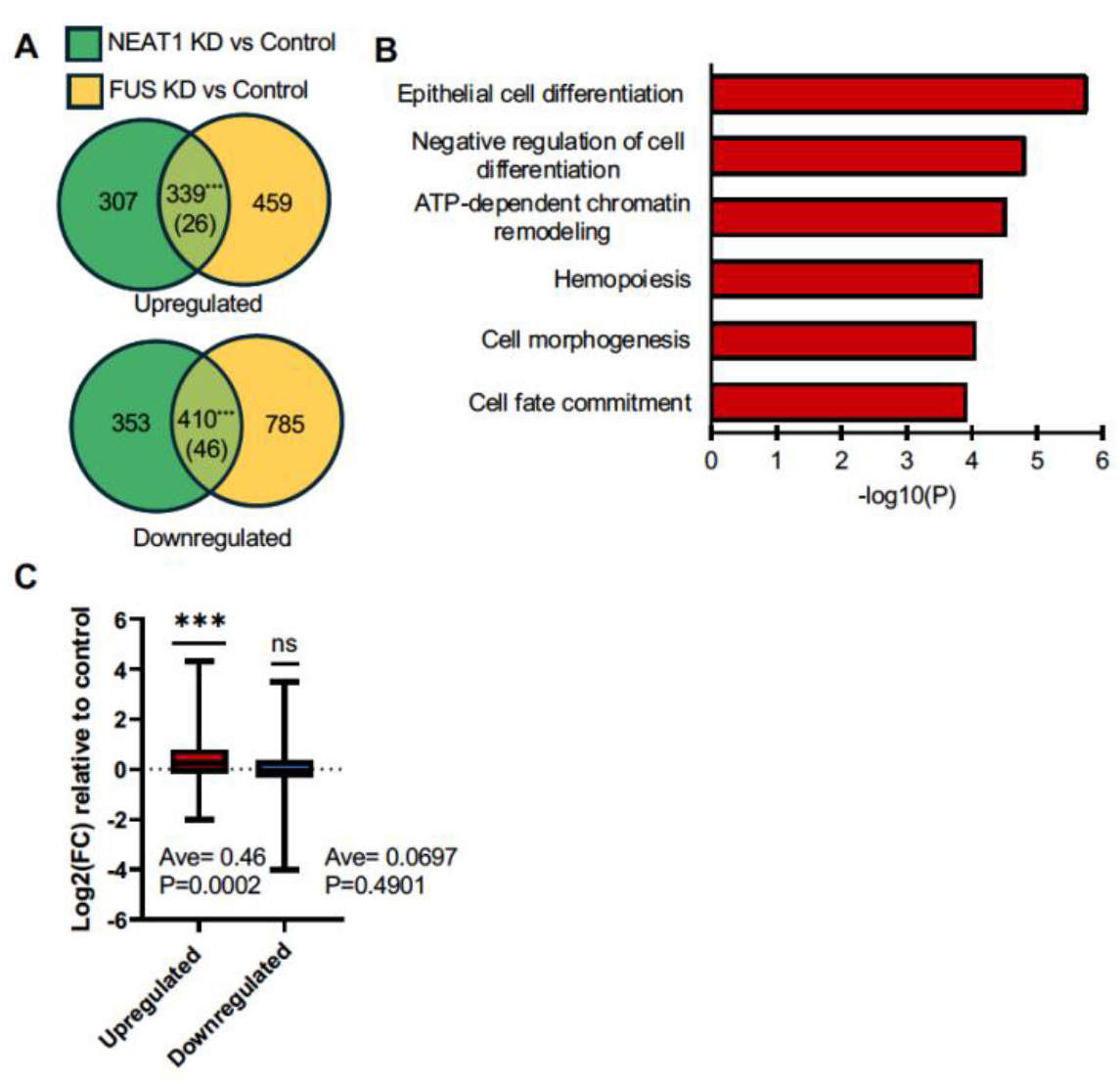
Paraspeckles regulate ERG mediated repression. (A) Overlap of differentially upregulated or downregulated genes in NEAT1 or FUS shRNA compared to scramble shRNA in VCaP cells. Number in parenthesis represents random expected overlap. *p*-values were calculated using fishers exact test, where ∗∗∗ indicates a *p*-value < 0.001 (B) Gene ontology of ERG relevant pathways from the upregulated paraspeckle signature. (C) Box-plots showing expression of paraspeckle signature genes ERG KD compared to control KD RNA-seq in VCaP cells. p-values were calculated using one sample t test, where ∗∗∗ indicates a p-value < 0.001

## Discussion

These findings indicate a role for the paraspeckle in Ewing sarcoma and ERG rearrangement positive prostate cancer through regulation of transcription. In representative cell lines, cancer-associated phenotypes were disrupted with paraspeckle loss. In Ewing sarcoma, paraspeckles and EWSR1::FLI1 promoted similar transcriptomes, and paraspeckle component SFPQ was found at the chromatin of direct EWSR1::FLI1 transcriptional targets. Further, SFPQ binding was disrupted when EWSR1::FLI1 was lost. Together, this supports a model that EWSR1::FLI1 can modulate paraspeckle genomic location, directing the paraspeckle to EWSR1::FLI1 targets to promote an oncogenic transcriptome.

There are several different ways the paraspeckle may be modulating EWSR1::FLI1 transcription at chromatin. One potential mechanism is that the paraspeckle is acting as a hub of transcriptional machinery that EWSR1::FLI1 can direct to target genes. Paraspeckles have been linked with active chromatin and contain important transcriptional machinery including Pol II as well as nascent mRNA (32, 36). Another potential mechanism involves modulation of regulatory DNA components. In Ewing sarcoma, EWSR1::FLI1 targets DNA repeats of GGAA, known as GGAA microsatellites (37, 38). These microsatellites have been identified as having either promoter or enhancer attributes, with qualities of both being dependent on distance from target genes and GGAA microsatellite length (37). The paraspeckle may serve as a connection between enhancer and promoter microsatellites that are genomically distant from each other or their target genes. Similarly, it has been shown that multiple copies of FLI1 can bind to GGAA microsatellites (39). Therefore, the paraspeckle may act as a concentrating agent, sequestering multiple copies of EWSR1::FLI1 to increase the number of EWSR1::FLI1 proteins bound to a GGAA microsatellite to modulate transcription.

The role of paraspeckles in ERG-positive prostate cancer is not as clear as in Ewing sarcoma, as ERG transcriptional repression was affected by paraspeckle disruption, but ERG associated transcriptional activation was not. However, it is not surprising that ERG and EWSR1::FLI1 would show different regulation by the paraspeckle. We have previously shown that ERG alone cannot rescue EWSR1::FLI1 knockdown in Ewing sarcoma cell lines (40) due to regulation by a co-repressor complex including EZH2. A mutant version of ERG that does not bind this repressive complex can rescue EWSR1::FLI1 knockdown (41). Unlike ERG, EWSR1::FLI1 does not bind this repressive complex. There is previous evidence of interplay between EZH2 and the paraspeckle or NEAT1 (42–44). Therefore, it is possible that ERG repression in prostate cancer may be modulated by the paraspeckle through EZH2. Loss of paraspeckles could then disrupt EZH2 repression of ERG, relieving repression of ERG target genes. Several genes that were upregulated with paraspeckle disruption in VCaP cells and linked to ERG related pathways are also regulated by EZH2, including *DAB2IP, RAP1GAP*, and *NKX2-2* (45–47).

Although our gene expression data in ERG-positive prostate cancer and Ewing sarcoma did not fully recapitulate each other, there was a consistent modulation of cell state with paraspeckle loss in both models. In Ewing sarcoma, our RNA-seq analysis and phenotypic assays showed that both paraspeckles and EWSR1::FLI1 promote pro-growth transcription factor pathways (E2F, MYC) and repress mesenchymal differentiation, consistent with the previously published role of EWSR1::FLI1 (29). In ERG-positive prostate cancer, our RNA-seq analysis showed that loss of paraspeckle function activated genes associated with epithelial differentiation and hematopoiesis. This is consistent with our Ewing sarcoma model with both models moving towards a more differentiated state with paraspeckle loss. The canonical roles of ERG and FLI1 are in maintenance of hematopoietic stem cells and the regulation of cell fate (34, 48). Similarly, there have been established roles of the paraspeckle in cell differentiation (36, 43). Therefore, disruption of paraspeckles is likely to promote movement to a more differentiated state in ETS positive cancers. This potentially limits the plasticity of cancer cells, preventing the ability of cells to respond to new environments, such as losing survivability in single cell state as reflected in soft agar and clonogenic growth assays.

As with most transcription factors, it has been difficult to develop drugs with the necessary specificity for ERG or FLI1. Our data indicate that disruption of paraspeckles has a significant effect on ERG positive prostate cancer and Ewing sarcoma cell survival in oncogenic phenotypic assays, but not a significant toxic effect on cell proliferation. This suggests that paraspeckles may be a good target for drug discovery in ETS-positive cancers. For example, proteins such as hnRNPK regulate paraspeckle formation by modulating NEAT1_2 transcription and already have available inhibitors (49). This study focused on the role of paraspeckles in ERG-positive prostate cancer and EWSR1::FLI1 dependent Ewing sarcoma. However, there are other cancers dependent on oncogenic ETS and EWSR1 interactions, including ETV4, ETV5, or ETV1-positive prostate cancer. There are also other cancers that are dependent on fusions of paraspeckle components and transcription factors, including SFPQ::TFE3 in renal cell carcinoma. The paraspeckle is a strong candidate for further mechanistic investigation and drug development.

### Experimental Procedures

#### Cell lines and immunoblots

A673 (ATCC, CRL-1598), VCaP (ATCC, CRL-2876), and HEK293t (ATCC, CRL-3216) cells were grown in Dulbecco’s modified Eagle’s media (DMEM; Corning). PC-3 cells (ATCC, CRL-1435) were grown in F-12K media (Corning). All media was supplemented with 10% fetal bovine serum (Sigma) and with 1× penicillin/streptomycin (Corning). Cell lines were authenticated using PowerPlex 16 HS assay with > 80% match to eight core short tandem repeat loci. Cell lines were tested for *mycoplasma* every four months using a mycoplasma PCR detection kit (Abcam). All cells were incubated at 37 °C and 5% CO_2_. All shRNAs were packaged in lentiviral vectors produced by cotransfecting HEK-293T cells with pMDLg/pRRE (Addgene plasmid 12251), pRSV-Rev (Addgene plasmid 12253), and pMD2.G (Addgene plasmid 12259), packaging plasmids in addition to the pLKO.1 cloning vector (Addgene plasmid 8453) with the corresponding shRNA sequence. Retroviral protein expression vectors were created using the methods described in (27). Cell extracts were run on SDS-page gels, transferred to nitrocellulose membrane, blocked with 5% milk TBS, incubated with 1:1000 primary antibody 5% milk TBS overnight, 1:12,500 secondary antibody 5% milk TBS 2 h, exposed to Femto ECL, and imaged on a Chemdoc (Biorad). Antibodies used were Flag (Millipore Sigma; SMR15), GAPDH (Santa Cruz Biotechnology; sc-47724), ERG (Biocare Medical; CM421C), and SFPQ (Abcam; Ab38148).

#### Droplet assay

Proteins were purified as previously described (27, 50). Fluorescent tags were cloned into pET28a vector (Novagen) and then used as the backbone for EWSR1and ERG cloning. Proteins were induced with 0.5 mM IPTG in BL21 *Escherichia coli* cells, grown for 3 h at 37 °C, and then purified with nickel-nitrilotriacetic acid beads (Qiagen). Proteins were diluted to specified concentration in droplet buffer (20 mM Na_2_HPO_4_/NaH_2_PO_4_, pH 7.5, 75 mM NaCl, 2.5% glycerol, 1 mM DTT) (51) and then centrifuged at 4°C. Proteins were mixed equimolar and/or treated with 10% of PEG (Sigma), 1,6 Hexanediol (Sigma), or 2,5 Hexanediol (Sigma). Mixture was placed on a glass slide, covered with a coverslip, and imaged at 100X on an Eclipse E600 (Nikon).

#### RNA-Immunoprecipitation

6×10^5^ cells were plated and incubated for 24 h at 37°C. Plates were washed with cold PBS (137 mM NaCl, 2.7 mM KCl, 10 mM Na_2_HPO_4_, and 1.8 mM KH_2_PO_4_) then fixed with UV Crosslinker (Stratagene) at 150 mj/cm^2^. Cells were resuspended in high salt buffer (20 mM Tris-HCl pH 7.5, 500 mM NaCl, 10 mM EDTA, 10% glycerol) and sonicated. Lysate was added to Anti-FLAG® M2 Magnetic Beads (Sigma) and incubated at 4°C for 2 h. Beads were treated with proteinase K (Sigma) and RNA was isolated with Trizol (ThermoFisher) following provider instructions. RNA levels were measured by reverse transcription followed by qRT-PCR with standard curves as described previously (52).

#### FISH and IF

1×10^4^ cells were plated on a coverslip in a 24 well plate. Cells were incubated at 37°C until around 80% confluency. Cells were fixed with 4% paraformaldehyde for 20 min, then permeabilized with 70% ethanol for 1 h, and then blocked with TBS (10 mM Tris, 150 mM NaCl, pH 7.4) containing 0.1% Tween-20, 1% BSA, and 10% horse serum for 1hr. Primary antibodies were diluted in TBS with 1% BSA and incubated with cells for 1 h at RT. Secondary antibody (ThermoFisher, A32790) diluted in the same buffer and incubated with cells for 1 h at RT. Coverslips were mounted onto glass slides using VECTASHIELD mounting media with 4′,6-diamidino-2-phenylindole (Vector Laboratories H-1200) and sealed with nail polish. RNA-FISH preparation was the same as IF until blocking. After permeabilization, slides were incubated for 5 min in 2X Saline-sodium citrate buffer (SSC) with 10% de-ionized formamide. Slides were then placed on top of hybridization buffer (10% de-ionized formamide, 2X SSC, 10% dextran sulfate, 0.1 mg/µl yeast tRNA) with Quasar 570 Dye conjugated Neat1 FISH Probe (Biosearch Technologies; SMF-2036-1) in a humidified chamber. Coverslips were incubated at 37°C for 16 hours. Coverslips were then washed with 4X SSC and mounted onto glass slide as with IF. If IF and RNA-FISH were performed together, IF was done first followed by a second fixation with 4% paraformaldehyde before proceeding to RNA-FISH.

#### Soft agar and clonogenic growth

For soft agar, 2 mL of 1% agar in DMEM was plated in a 6 well plate. Once the first layer was solidified, 1000 cells suspended in 1 mL of 0.35% agar and media were plated. Once solidified, a feeding layer of 2 mL of media was added. Cells were incubated at 37°C until 3 weeks. The wells were imaged using UV and analyzed through measuring the area of a thresholded image using ImageJ. For clonogenic growth, 1×10^4^ cells were plated in 2 mL of DMEM and incubated at 37°C for 2 weeks. The cells were fixed with 10% formalin and stained with crystal violet. Images of the wells were taken and analyzed the same way as soft agar.

#### Cell migration

Trans*-*well migration assay was performed as previously described (53). Briefly, 8 μM pore size Boyden chambers (VRW) were placed in a 24-well plate filled with 750 μl of DMEM. 5×10^4^ cells suspended in 100 μl of DMEM without FBS were plated into a Boyden chamber and incubated at 37°C for 48 h. The membranes were stained with Hema 3 staining kit (ThermoFisher) and dried for 24 h before being imaged and analyzed through measuring the area of a thresholded image using ImageJ. Each membrane was imaged 5 times, and the average of those areas were normalized to control.

#### Cell proliferation

Cells were plated in 100 µL of media in a 96 well plate, 2000 cells for A673 cells and 5000 cells for VCaP cells, and incubated at 37°C for indicated times. 5 mg/mL of 3-(4,5-Dimethylthiazol-2-yl)-2,5-Diphenyltetrazolium Bromide (MTT) was added to the cells and were incubated at 37°C for 4 h. Media was aspirated and 100 µL of DMSO was added. Absorbance at 570 nm was measured. Day 3 was normalized to day 1 before being normalized to control. For TC71 proliferation, cells were transduced with viral vectors for 48 h before following the same protocol as the A673 cells above. Absorbance was collected for 4 days and each day was normalized to day 1 before being plotted.

#### RNA-seq

RNA was isolated from cells with Trizol (ThermoFisher) following provider instructions. Isolated RNA was sent for sequencing on the Illumina NovaSeq X Plus platform through Plasmidsaurus. Analysis on provided fastq files was performed through galaxy (https://galaxyproject.org). Alignments and counts were performed through HISAT2 and featureCounts. Differential gene expression was determined using DESeq2 with an adjp-value cut off of <0.05 and a |log2(FC)| cut off of >1. A673 cells were done in technical triplicate and VCaP cells were done in technical duplicate. For A673 cells, pathway enrichment was done through GSEA while VCaP cells gene ontology analysis was done through metascape (https://metascape.org).

#### CHIP

ChIP was performed as previously described (24). Briefly, Dynabeads™ M-280 Sheep Anti-Rabbit IgG (ThermoFisher) were diluted in dilution buffer (20 mM Tris pH 7.9, 2 mM EDTA, 150 mM NaCl, 1% Triton X-100, 2 mg/ml BSA) and mixed with 5 uL SFPQ antibody (Ab38148, Abcam) and rotated overnight at 4°C. Cells were fixed using 1% formaldehyde (ThermoFisher) and quenched with 2 M glycine. Cells were lysed and sonicated (23) and lysate was added to beads for 4 hr at 4°C. Beads and input were treated with RNAase A (ThermoFisher) and proteinase K (Sigma) before being isolated through phenol/chloroform (Sigma) extraction. Target DNA was quantified through qPCR as in RIP.

## Data availability

RNA-seq data are available at Gene Expression Omnibus (www.ncbi.nlm.nih.gov/geo/) under accession number GSE350110.

## Supporting Information

This article contains supporting information.

## Acknowledgements

The authors would like to thank Didier Trono (addgene #12251, 12253, 12259) and Bob Weinberg (addgene #8453) for plasmids.

## Funding and additional information

R.A.K. was supported by an NIH training grant (T32GM131994). The content of this paper is solely the responsibility of the authors and does not necessarily represent the official views of the National Institutes of Health.

## References

1. Tomlins, S. A., Laxman, B., Dhanasekaran, S. M., Helgeson, B. E., Cao, X., Morris, D. S. et al. (2007) Distinct classes of chromosomal rearrangements create oncogenic ETS gene fusions in prostate cancer Nature 448, 595–599 nature06024 [pii]10.1038/nature06024

2. Helgeson, B. E., Tomlins, S. A., Shah, N., Laxman, B., Cao, Q., Prensner, J. R. et al. (2008) Characterization of TMPRSS2:ETV5 and SLC45A3:ETV5 gene fusions in prostate cancer Cancer Res 68, 73–80 68/1/73 [pii]10.1158/0008-5472.CAN-07-5352

3. Grunewald, T. G. P., Cidre-Aranaz, F., Surdez, D., Tomazou, E. M., de Alava, E., Kovar, H., et al. (2018) Ewing sarcoma Nat Rev Dis Primers 4, 5 10.1038/s41572-018-0003-x

4. Kedage, V., Selvaraj, N., Nicholas, T. R., Budka, J. A., Plotnik, J. P., Jerde, T. J. et al. (2016) An Interaction with Ewing’s Sarcoma Breakpoint Protein EWS Defines a Specific Oncogenic Mechanism of ETS Factors Rearranged in Prostate Cancer Cell Rep 17, 1289–1301 10.1016/j.celrep.2016.10.001

5. Schwartz, J. C., Cech, T. R., and Parker, R. R. (2015) Biochemical Properties and Biological Functions of FET Proteins Annu Rev Biochem 84, 355–379 10.1146/annurev-biochem-060614-034325

6. Ahn, J. H., Davis, E. S., Daugird, T. A., Zhao, S., Quiroga, I. Y., Uryu, H., et al. (2021) Phase separation drives aberrant chromatin looping and cancer development Nature 595, 591-+ 10.1038/s41586-021-03662-5

7. Sabari, B. R., Dall’Agnese, A., Boija, A., Klein, I. A., Coffey, E. L., Shrinivas, K., et al. (2018) Coactivator condensation at super-enhancers links phase separation and gene control Science 361, 10.1126/science.aar3958

8. Boulay, G., Sandoval, G. J., Riggi, N., Iyer, S., Buisson, R., Naigles, B. et al. (2017) Cancer-Specific Retargeting of BAF Complexes by a Prion-like Domain Cell 171, 163–178 e119 10.1016/j.cell.2017.07.036

9. Clemson, C. M., Hutchinson, J. N., Sara, S. A., Ensminger, A. W., Fox, A. H., Chess, A. et al. (2009) An architectural role for a nuclear noncoding RNA: NEAT1 RNA is essential for the structure of paraspeckles Molecular cell 33, 717–726 10.1016/j.molcel.2009.01.026

10. West, J. A., Mito, M., Kurosaka, S., Takumi, T., Tanegashima, C., Chujo, T. et al. (2016) Structural, super-resolution microscopy analysis of paraspeckle nuclear body organization J Cell Biol 214, 817–830 10.1083/jcb.201601071

11. Fox, A. H., Bond, C. S., and Lamond, A. I. (2005) P54nrb forms a heterodimer with PSP1 that localizes to paraspeckles in an RNA-dependent manner Mol Biol Cell 16, 5304–5315 10.1091/mbc.e05-06-0587

12. Hirose, T., Yamazaki, T., and Nakagawa, S. (2019) Molecular anatomy of the architectural NEAT1 noncoding RNA: The domains, interactors, and biogenesis pathway required to build phase-separated nuclear paraspeckles Wiley Interdiscip Rev RNA 10, e1545 10.1002/wrna.1545

13. Li, R., Harvey, A. R., Hodgetts, S. I., and Fox, A. H. (2017) Functional dissection of NEAT1 using genome editing reveals substantial localization of the NEAT1_1 isoform outside paraspeckles RNA 23, 872–881 10.1261/rna.059477.116

14. Chakravarty, D., Sboner, A., Nair, S. S., Giannopoulou, E., Li, R., Hennig, S., et al. (2014) The oestrogen receptor alpha-regulated lncRNA NEAT1 is a critical modulator of prostate cancer Nature communications 5, 5383 10.1038/ncomms6383

15. Hirose, T., Virnicchi, G., Tanigawa, A., Naganuma, T., Li, R., Kimura, H. et al. (2014) NEAT1 long noncoding RNA regulates transcription via protein sequestration within subnuclear bodies Mol Biol Cell 25, 169–183 10.1091/mbc.E13-09-0558

16. Prasanth, K. V., Prasanth, S. G., Xuan, Z., Hearn, S., Freier, S. M., Bennett, C. F. et al. (2005) Regulating gene expression through RNA nuclear retention Cell 123, 249–263 10.1016/j.cell.2005.08.033

17. Yu, X., Li, Z., Zheng, H., Chan, M. T., and Wu, W. K. (2017) NEAT1: A novel cancer-related long non-coding RNA Cell Prolif 50, 10.1111/cpr.12329

18. Zeng, C., Xu, Y., Xu, L., Yu, X., Cheng, J., Yang, L., et al. (2014) Inhibition of long non-coding RNA NEAT1 impairs myeloid differentiation in acute promyelocytic leukemia cells BMC Cancer 14, 693 10.1186/1471-2407-14-693

19. Choudhry, H., Albukhari, A., Morotti, M., Haider, S., Moralli, D., Smythies, J. et al. (2015) Tumor hypoxia induces nuclear paraspeckle formation through HIF-2alpha dependent transcriptional activation of NEAT1 leading to cancer cell survival Oncogene 34, 4482–4490 10.1038/onc.2014.378

20. Bond, C. S., and Fox, A. H. (2009) Paraspeckles: nuclear bodies built on long noncoding RNA J Cell Biol 186, 637–644 10.1083/jcb.200906113

21. Nitusca, D., Marcu, A., Dema, A., Balacescu, L., Balacescu, O., Bardan, R., et al. (2021) Long Noncoding RNA NEAT1 as a Potential Candidate Biomarker for Prostate Cancer Life (Basel) 11, 10.3390/life11040320

22. Kauffman, E. C., Ricketts, C. J., Rais-Bahrami, S., Yang, Y., Merino, M. J., Bottaro, D. P. et al. (2014) Molecular genetics and cellular features of TFE3 and TFEB fusion kidney cancers Nat Rev Urol 11, 465–475 10.1038/nrurol.2014.162

23. Kwon, I., Kato, M., Xiang, S., Wu, L., Theodoropoulos, P., Mirzaei, H. et al. (2013) Phosphorylation-regulated binding of RNA polymerase II to fibrous polymers of low-complexity domains Cell 155, 1049–1060 10.1016/j.cell.2013.10.033

24. Greulich, B. M., Rajendran, S., Downing, N. F., Nicholas, T. R., and Hollenhorst, P. C. (2023) A complex with poly(A)-binding protein and EWS facilitates the transcriptional function of oncogenic ETS transcription factors in prostate cells The Journal of biological chemistry 299, 105453 10.1016/j.jbc.2023.105453

25. Tulpule, A., Guan, J., Neel, D. S., Allegakoen, H. R., Lin, Y. P., Brown, D. et al. (2021) Kinase-mediated RAS signaling via membraneless cytoplasmic protein granules Cell 184, 2649–2664 e2618 10.1016/j.cell.2021.03.031

26. Zheng, T., Wake, N., Weng, S. L., Perdikari, T. M., Murthy, A. C., Mittal, J. et al. (2025) Molecular insights into the effect of 1,6-hexanediol on FUS phase separation EMBO J 44, 2725–2740 10.1038/s44318-025-00431-2

27. Kedage, V., Selvaraj, N., Nicholas, T. R., Budka, J. A., Plotnik, J. P., Jerde, T. J. et al. (2016) An Interaction with Ewing’s Sarcoma Breakpoint Protein EWS Defines a Specific Oncogenic Mechanism of ETS Factors Rearranged in Prostate Cancer Cell Rep 17, 1289–1301 10.1016/j.celrep.2016.10.001

28. Naganuma, T., Nakagawa, S., Tanigawa, A., Sasaki, Y. F., Goshima, N., and Hirose, T. (2012) Alternative 3’-end processing of long noncoding RNA initiates construction of nuclear paraspeckles EMBO J 31, 4020–4034 10.1038/emboj.2012.251

29. Chaturvedi, A., Hoffman, L. M., Welm, A. L., Lessnick, S. L., and Beckerle, M. C. (2012) The EWS/FLI Oncogene Drives Changes in Cellular Morphology, Adhesion, and Migration in Ewing Sarcoma Genes Cancer 3, 102–116 10.1177/1947601912457024

30. Sole, A., Grossetete, S., Heintze, M., Babin, L., Zaidi, S., Revy, P. et al. (2021) Unraveling Ewing Sarcoma Tumorigenesis Originating from Patient-Derived Mesenchymal Stem Cells Cancer Res 81, 4994–5006 10.1158/0008-5472.CAN-20-3837

31. Riggi, N., Knoechel, B., Gillespie, S. M., Rheinbay, E., Boulay, G., Suva, M. L. et al. (2014) EWS-FLI1 utilizes divergent chromatin remodeling mechanisms to directly activate or repress enhancer elements in Ewing sarcoma Cancer Cell 26, 668–681 10.1016/j.ccell.2014.10.004

32. West, J. A., Davis, C. P., Sunwoo, H., Simon, M. D., Sadreyev, R. I., Wang, P. I. et al. (2014) The long noncoding RNAs NEAT1 and MALAT1 bind active chromatin sites Molecular cell 55, 791–802 10.1016/j.molcel.2014.07.012

33. Knudsen, K. J., Rehn, M., Hasemann, M. S., Rapin, N., Bagger, F. O., Ohlsson, E. et al. (2015) ERG promotes the maintenance of hematopoietic stem cells by restricting their differentiation Genes Dev 29, 1915–1929 10.1101/gad.268409.115

34. Taoudi, S., Bee, T., Hilton, A., Knezevic, K., Scott, J., Willson, T. A. et al. (2011) ERG dependence distinguishes developmental control of hematopoietic stem cell maintenance from hematopoietic specification Genes & development 25, 251–262 10.1101/gad.2009211

35. Sandoval, G. J., Pulice, J. L., Pakula, H., Schenone, M., Takeda, D. Y., Pop, M. et al. (2018) Binding of TMPRSS2-ERG to BAF Chromatin Remodeling Complexes Mediates Prostate Oncogenesis Molecular cell 71, 554–566 e557 10.1016/j.molcel.2018.06.040

36. Grosch, M., Ittermann, S., Rusha, E., Greisle, T., Ori, C., Truong, D. J., et al. (2020) Nucleus size and DNA accessibility are linked to the regulation of paraspeckle formation in cellular differentiation BMC Biol 18, 42 10.1186/s12915-020-00770-y

37. Johnson, K. M., Taslim, C., Saund, R. S., and Lessnick, S. L. (2017) Identification of two types of GGAA-microsatellites and their roles in EWS/FLI binding and gene regulation in Ewing sarcoma PLoS One 12, e0186275 10.1371/journal.pone.0186275

38. Guillon, N., Tirode, F., Boeva, V., Zynovyev, A., Barillot, E., and Delattre, O. (2009) The oncogenic EWS-FLI1 protein binds in vivo GGAA microsatellite sequences with potential transcriptional activation function PloS one 4, e4932 10.1371/journal.pone.0004932

39. Hou, C., and Tsodikov, O. V. (2025) Structure and cooperative formation of a FLI1 filament on contiguous GGAA DNA sites Nucleic Acids Res 53, 10.1093/nar/gkaf205

40. Downing, N. F., Mills, K. M., and Hollenhorst, P. C. (2026) PRC2/FOXO1-Mediated Repression Determines Interchangeability of ETS Oncogenes in Prostate Cancer and Ewing Sarcoma Molecular cancer research : MCR 24, 48–59 10.1158/1541-7786.MCR-25-0389

41. Kedage, V., Strittmatter, B. G., Dausinas, P. B., and Hollenhorst, P. C. (2017) Phosphorylation of the oncogenic transcription factor ERG in prostate cells dissociates polycomb repressive complex 2, allowing target gene activation The Journal of biological chemistry 292, 17225–17235 10.1074/jbc.M117.796458

42. Li, K., and Wang, Z. (2021) Speckles and paraspeckles coordinate to regulate HSV-1 genes transcription Commun Biol 4, 1207 10.1038/s42003-021-02742-6

43. Yin, C., Ge, Z., Yuan, J., Chen, Y., Tang, Y., Xiang, Y., et al. (2024) NEAT1 regulates VSMC differentiation and calcification in as long noncoding RNA NEAT1 enhances phenotypic and osteogenic switching of vascular smooth muscle cells in atherosclerosis via scaffolding EZH2 Am J Physiol Cell Physiol 326, C1721-C1734 10.1152/ajpcell.00587.2023

44. Wang, S., Zuo, H., Jin, J., Lv, W., Xu, Z., Fan, Y., et al. (2019) Long noncoding RNA Neat1 modulates myogenesis by recruiting Ezh2 Cell Death Dis 10, 505 10.1038/s41419-019-1742-7

45. Chen, H., Tu, S. W., and Hsieh, J. T. (2005) Down-regulation of human DAB2IP gene expression mediated by polycomb Ezh2 complex and histone deacetylase in prostate cancer J Biol Chem 280, 22437–22444 10.1074/jbc.M501379200

46. Banerjee, R., Mani, R. S., Russo, N., Scanlon, C. S., Tsodikov, A., Jing, X. et al. (2011) The tumor suppressor gene rap1GAP is silenced by miR-101-mediated EZH2 overexpression in invasive squamous cell carcinoma Oncogene 30, 4339–4349 10.1038/onc.2011.141

47. Landeira, D., Sauer, S., Poot, R., Dvorkina, M., Mazzarella, L., Jorgensen, H. F. et al. (2010) Jarid2 is a PRC2 component in embryonic stem cells required for multi-lineage differentiation and recruitment of PRC1 and RNA Polymerase II to developmental regulators Nat Cell Biol 12, 618–624 10.1038/ncb2065

48. Athanasiou, M., Mavrothalassitis, G., Sun-Hoffman, L., and Blair, D. G. (2000) FLI-1 is a suppressor of erythroid differentiation in human hematopoietic cells Leukemia 14, 439–445 10.1038/sj.leu.2401689

49. Shu, B., Zeng, P., Kang, S., Li, P. H., Hu, D., Kuang, G. et al. (2019) Syntheses and evaluation of new Quinoline derivatives for inhibition of hnRNP K in regulating oncogene c-myc transcription Bioorg Chem 85, 1–17 10.1016/j.bioorg.2018.12.020

50. Selvaraj, N., Kedage, V., and Hollenhorst, P. C. (2015) Comparison of MAPK specificity across the ETS transcription factor family identifies a high-affinity ERK interaction required for ERG function in prostate cells Cell Commun Signal 13, 12 10.1186/s12964-015-0089-7

51. Hofweber, M., Hutten, S., Bourgeois, B., Spreitzer, E., Niedner-Boblenz, A., Schifferer, M. et al. (2018) Phase Separation of FUS Is Suppressed by Its Nuclear Import Receptor and Arginine Methylation Cell 173, 706–719 e713 10.1016/j.cell.2018.03.004

52. Greulich, B. M., Plotnik, J. P., Jerde, T. J., and Hollenhorst, P. C. (2021) Toll-like receptor 4 signaling activates ERG function in prostate cancer and provides a therapeutic target NAR Cancer 3, zcaa046 10.1093/narcan/zcaa046

53. Hollenhorst, P. C., Ferris, M. W., Hull, M. A., Chae, H., Kim, S., and Graves, B. J. (2011) Oncogenic ETS proteins mimic activated RAS/MAPK signaling in prostate cells Genes & development 25, 2147–2157 10.1101/gad.17546311

