## Supplemental Files for "Oncogenic ETS factors expressed from fusion genes utilize EWSR1 to associate with the paraspeckle and regulate transcription"

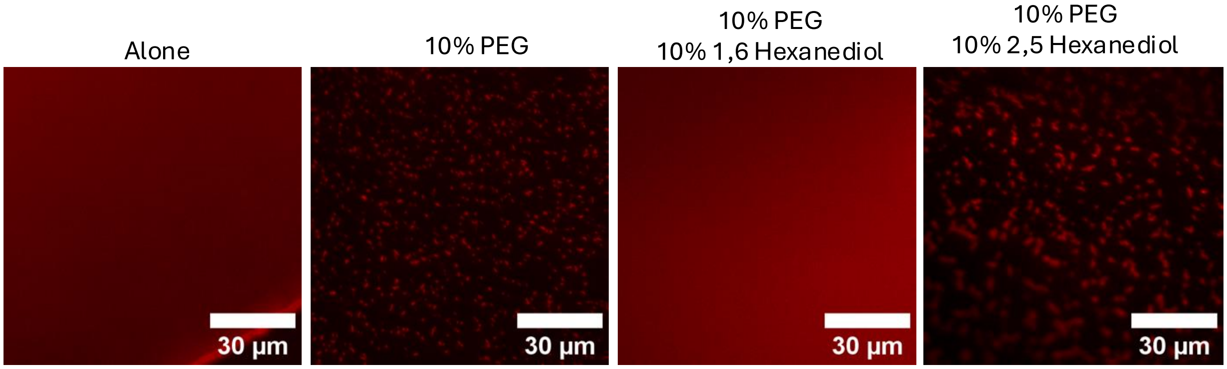


**Supporting Information Figure S1. EWSR1 droplet formation is reversible.** 3µM EWS_mCherry was combined with 10% PEG alone or with either 10% 1,6 hexanediol or 10% 2,5 hexanediol.

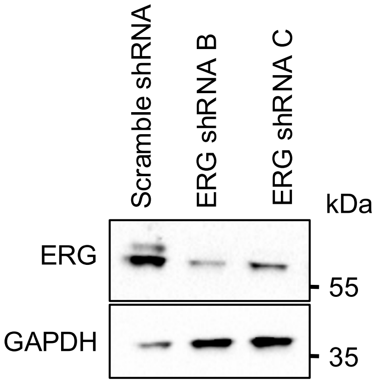


**Supporting Information Figure S2. ERG knockdown in VCaP cells**
Immunoblot from VCaP cells expressing indicated shRNAs.

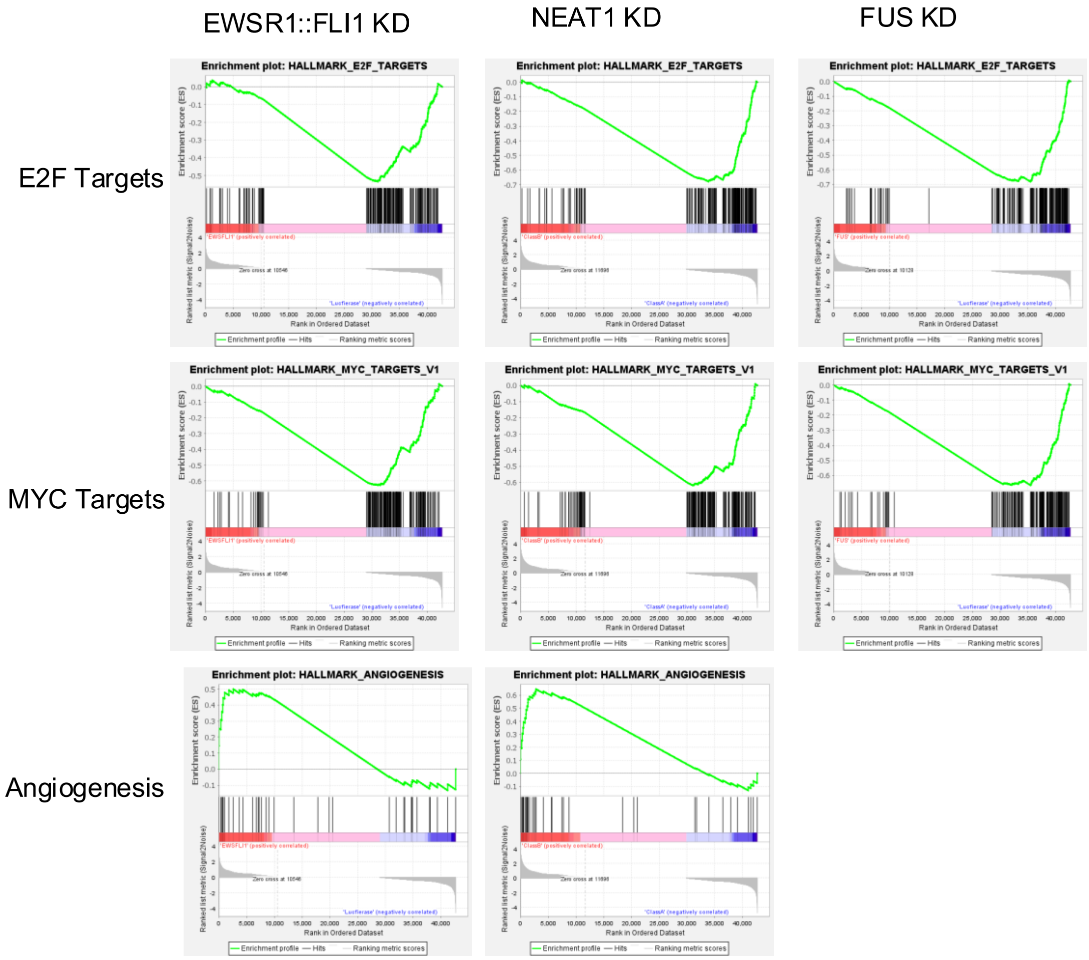


**Supporting Information Figure S3. EWSR1::FLI1 and paraspeckle knockdowns share trends in pathway enrichment of hallmark cancer pathways**
Gene set enrichment analysis of EWSR1::FLI1 shRNA, NEAT1 shRNA, and FUS shRNA each compared to luciferase shRNA in A673 cells.

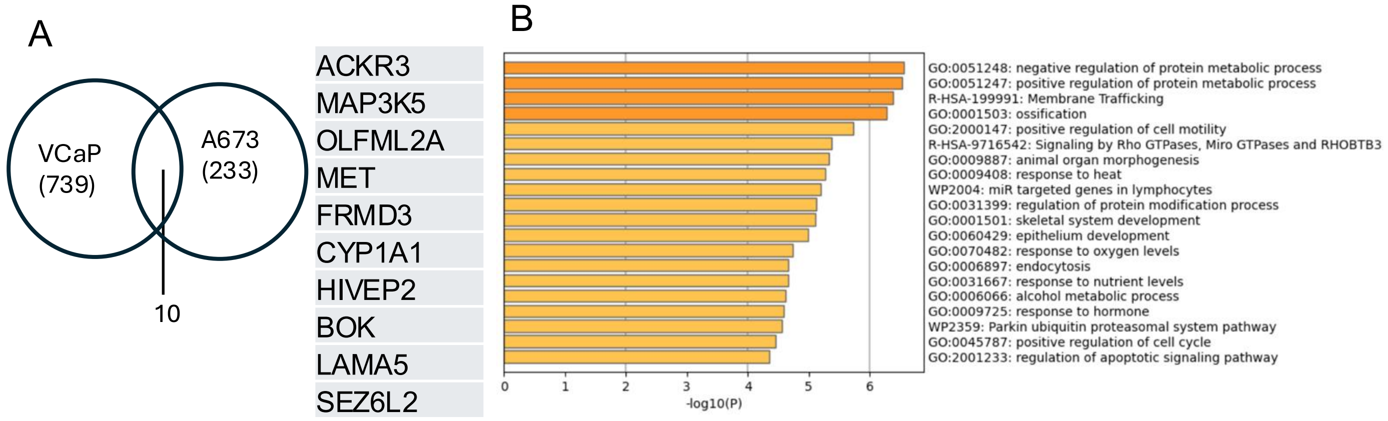


**Supporting Information Figure S4. Related to VCaP RNA-seq with NEAT1 and FUS knockdown.**

(A) Overlap between paraspeckle signature of A673 cells and VCaP cells. Ten genes in overlap are shown. (B) Most overrepresented gene ontologies for downregulated genes in both NEAT1 and FUS shRNA compared to scramble shRNA in VCaP cells.

Supporting Table S1. Target Sequences and Primers

| | Forward | Reverse |
|---|---|---|
| NEAT1 shRNA A | AAATGCAGGTTGAAGGGAATT | |
| FUS shRNA A | AACGATTATACCCAACAAGCA | |
| NEAT1_2 A shRNA | CACAGATAGAACTGGTATTAT | |
| NEAT1_2 B shRNA | AATTCAGAAGACCTTGAGGGC | |
| FUS shRNA B | AACACAGGCTATGGAACTCAGT | |
| ERG shRNA A | AAGGAACTCTCCTGATGAATG | |
| ERG shRNA B | GGCCAGATTTACCATATGA | |
| ERG shRNA C | CCCAAATACTTCAGTATAT | |
| NEAT1_2 qPCR | TGCGGGGATCTCTTTTGCAT | CAGATGTGCCAAGAGCCAGA |
| FUS qPCR | CCCTAAACCAGATGGCCCAG | CGATCATAGCCTCCTCTGCC |
| EWS/FLI1 qPCR | AAC CGA GCA GCT ATG GAC AGC | CTC CAG GAG GAA TTG CCA CA |
| RBP1 chr3: 139539447-139539742 | GCAGGTACTCCTCGAAATTCTC | CACTCCCCGAAATGCCAG |
| SLIT2 chr4: 19973622-19973811 | CTCTAGAACACATTGCAAGGATG | TCTCATCCCTCTTCCCTCC |
| TNC chr9: 115369631-115369847 | GCATTCAATCCTTGATTCCCC | AGTTCGCCAAAGCCCTAAAG |
| ANKRD7 chr7: 118311550-118311788 | GGAAATGAGTATGAGTTGTCTATTGC | CACTACACTGAATCTAGGAAGCA |
